# A genetic interaction screen in *Streptococcus pneumoniae* identifies functionally redundant vaccine candidate proteins CbpC and CbpJ

**DOI:** 10.1101/2021.04.07.438901

**Authors:** Lisa M. Russo, Allison J. Matthews, Revati F. Masilamani, David W. Lazinski, Andrew Camilli

**Author notes:** Address correspondence to Andrew Camilli. These authors contributed equally to this study. LMR is the first author due to the lead role in experimentation and analysis.

## Abstract

*Streptococcus pneumoniae* is a Gram-positive bacterium that asymptomatically colonizes the nasopharynx and can disseminate to sterile sites resulting in pneumococcal diseases such as pneumonia, otitis media, bacteremia, and meningitis. Due to increased incidence of invasive disease caused by serotypes that are not included in available polysaccharide vaccines, there is a need for a broadly protective protein vaccine to complement the polysaccharide based vaccines. To limit immune escape such a vaccine would ideally target proteins that are essential for virulence. However, the genetic robustness of *S. pneumoniae* results in few surface exposed proteins being essential for virulence. Here we carried out a genetic interaction screen to identify functionally redundant surface protein pairs that could be used as bivalent protein vaccines, based on the observation that together, these protein pairs are essential for virulence. We identified four pairs of functionally redundant surface proteins that displayed a significant competitive disadvantage during murine pneumococcal pneumonia. Immunization with the most attenuated pair, CbpC and CbpJ, resulted in production of high titers of specific antibodies and a modest increased median survival times of mice challenged with pneumococcal pneumonia. This study demonstrates a method to identify essential pairs of surface-associated virulence proteins that could be widely applied to many bacterial pathogens.

**IMPORTANCE:** Infection by *Streptococcus pneumoniae* can result in life-threatening illness. Current licensed polysaccharide vaccines only protect against serotypes that are present in the vaccine – at most 23 serotypes of the total 100 identified, circulating serotypes. There remains a need for a widely protective protein vaccine that is effective against most circulating strains of *S. pneumoniae*. The significance of our research is in developing a method to identify functionally redundant protein pairs as potential vaccine candidates, which could inform the development of effective bivalent protein vaccines for many bacterial pathogens.

## INTRODUCTION

*Streptococcus pneumoniae* (the pneumoccoccus) remains a serious international public health concern. *S. pneumoniae* is a Gram-positive bacterium that asymptomatically colonizes the upper respiratory tract of significant portions of the population (1–3). Dissemination of bacteria to sterile sites throughout the body can result in life threatening diseases such as pneumonia, bacteremia, and meningitis. Globally, *S. pneumoniae* is the leading cause of community acquired pneumonia and lower respiratory morbidity and mortality (4–7). Increasing levels of antibiotic resistance have made this pathogen more challenging and more expensive to treat, making widespread vaccination critical to reduce the public health burden resulting from pneumococcal infection (8–11).

Existing *S. pneumoniae* vaccine strategies target the bacterial capsule, a layer of polysaccharide that coats the cell surface and distinguishes between the 100 distinct serotypes (12). Currently available vaccines protect against the serotypes of *S. pneumoniae* that are responsible for the majority of invasive pneumococcal disease (1). Two major vaccines are commonly used: a 13-valent pneumococcal conjugate vaccine (PCV13, Prevnar 13) and a 23-valent pneumococcal polysaccharide vaccine (PPSV23, Pneumovax 23). Initial introduction of the conjugate vaccine (PCV7) resulted in a significant decrease in invasive pneumococcal disease (13). However, these vaccines are only protective against serotypes represented in the vaccine. Over time widespread immunization in a population can result in increased frequency of disease from non-vaccine serotypes, a phenomenon known as serotype replacement (14–20). In addition, *S. pneumoniae* are able to undergo homologous recombinations that exchange capsular loci, resulting in serotype switching (14, 21). Serotype replacement and switching are forms of immune evasion that allow the bacteria to evade adaptive immune responses elicited by vaccination. This limits the efficacy of polysaccharide vaccines in the long term, requiring the development of vaccines with progressively increasing numbers of polysaccharide types to include emerging serotypes.

Therefore, there is a need for a broadly protective protein vaccine unaffected by serotype replacement and switching. We propose that one or more well conserved proteins from *S. pneumoniae* that are essential for virulence and are accessible to the immune system could be an effective solution that could be used alone or in combination with the polysaccharide-based vaccines. Use of essential virulence proteins as vaccine antigens would serve to limit the potential for immune evasion by *S. pneumoniae*.

In this study, we demonstrate a new method to identify candidate proteins for pneumococcal vaccine development. We used Tn-seq data to identify singly essential surface proteins as well as genetic interaction screening of surface proteins to identify sets of functionally redundant gene pairs that, when deleted, severely attenuate nasopharynx colonization and lung infection. Immunization with the most attenuating pair, CbpC and CbpJ, elicited production of specific antibodies and extended survival in a mouse model of pneumococcal pneumonia, thereby indicating the feasibility of this strategy and the potential of this method to identify candidate vaccine antigens against other pathogens.

## RESULTS

### Selection of pneumococcal surface proteins as vaccine candidates

In an effort to identify potential protein vaccine candidates against *S. pneumoniae*, we initially chose to focus on single proteins that are both surface exposed and therefore more accessible to the immune system, and essential for viability in order to prevent immune evasion by the bacterium. We utilized previously published Tn-seq data (22) in serotype 4 strain TIGR4 to identify five surface proteins that are putatively essential for viability (Table S1). However, when tested as monovalent vaccines, none of these proteins were protective against pneumococcal lung infection (Fig. S1).

As an alternative approach, we chose to use high-throughput screening techniques to identify functionally redundant surface protein pairs as vaccine candidates. Previous high throughput mutant screens have demonstrated that the majority of *S. pneumoniae* surface proteins are dispensable for viability and virulence (22–25). We propose that there may be functional redundancy amongst some of these surface proteins. Therefore, our second approach aimed to identify a functionally redundant pair of surface proteins, which by themselves are not essential for virulence, but in combination are indispensable. A bivalent vaccine targeting such a functionally redundant pair of cell surface proteins could provide protection against immune evasion.

In order to identify functionally redundant pairs that could be used as bivalent vaccine antigens, we selected 28 *S. pneumoniae* candidate surface proteins for genetic interaction screening based on the following criteria: (1) They are predicted to be surface localized based on conserved domains (LPXTG motif, cell wall binding domain, peptidoglycan binding domain, and choline binding protein) (26–29), (2) The extracellular domain of each protein is well conserved across most sequenced strains of *S. pneumoniae* (88-100% amino acid identity), (3) The protein is dispensable for virulence in both nasopharyngeal colonization and lung infection (22), (4) The protein is not well-conserved in other bacterial or human genomes, and (5) Proteins previously demonstrated to be immunogenic in humans or mice were prioritized (30–34). These selected candidates are listed in Table 1.

**Table 1.** Non-essential surface protein candidates selected for genetic interaction screening

| Locus<sup>a</sup> | Gene | Domain(s)<sup>b</sup> | Name/Function | Fitness<sup>c</sup> |  |
| --- | --- | --- | --- | --- | --- |
|  |  |  |  | Naso | Lung |
| SP_0057 | *strH* | LPXTG | Beta-N-acetylhexosaminidase | 1 | 1 |
| SP_0069 | *cbpl* | CBP | Choline binding protein I | 1.1 | 0.9 |
| SP_0071 | *zmpC* |  | Zinc metalloprotease | 1 | 1 |
| SP_0082 |  | LPXTG | Cell wall surface anchor family protein | 1 | 1 |
| SP_0107 |  | PBD | LysM domain-containing protein | <0.1 | 0.7 |
| SP_0117 | *pspA* | CBP | Pneumococcal surface protein A | 1.1 | 0.9 |
| SP_0268 |  | LPXTG | Putative alkaline amylopullulanase | 1 | 0.9 |
| SP_0314 |  | LPXTG | Hyaluronidase | 1.1 | 1 |
| SP_0368 |  | LPXTG | Cell wall surface anchor family protein | 1 | 1 |
| SP_0377 | *cbpC* | CBP | Choline binding protein C | 1.1 | 1 |
| SP_0378 | *cbpJ* | CBP | Choline binding protein J | 1.1 | 0.8 |
| SP_0390 | *cbpG* | CW, CBP | Choline binding protein G | 1 | 0.9 |
| SP_0498 |  | LPXTG | Endo-beta-N-acetylglucosaminidase | 0.9 | 1 |
| SP_0641 |  | LPXTG | Serine protease, subtilase family | 1 | 1 |
| SP_0648 | *bgaA* | LPXTG | Beta-galactosidase | 1 | 1 |
| SP_0664 | *zmpB* |  | Zinc metalloprotease | 1 | 1 |
| SP_0667 |  | CBP | Putative surface protein | 0.6 | 1 |
| SP_0930 | *cbpE* | CBP | Choline binding protein E | 0.4 | 0.8 |
| SP_1154 | *iga* |  | IgA zinc metalloprotease | 1.1 | 1 |
| SP_1573 | *lytC* | CBP | lysozyme | 0.2 | 1 |
| SP_1772 |  | LPXTG | Cell wall surface anchor family protein | <0.1 | 0.6 |
| SP_1923 | *ply* |  | Pneumolysin | 0.3 | 0.9 |
| SP_1937 | *lytA* | CBP, CW | Autolysin | 0.2 | 0.7 |
| SP_1992 |  | LPXTG | Cell wall surface anchor family protein | <0.1 | 0.7 |
| SP_2136 | *pcpA* | CBP | Choline binding protein A | 0.8 | 1 |
| SP_2145 |  | LPXTG | Antigen, cell wall anchor family | 1 | 1 |
| SP_2190 | *cbpA* | CBP, CW | Choline binding protein A | 0.7 | 1 |
| SP_2201 | *cbpD* | CBP | Choline binding protein D | 1.1 | 0.9 |

a. Locus in strain TIGR4
b. Peptidoglycan binding domain (PBD), Cell wall binding domain (CW), Choline binding protein (CBP), Sortase-mediated cell wall anchoring motif (LPXTG)
c. Fitness values were rounded from citation (van Opijnen & Camilli, 2012). A fitness of 1 indicates that the mutant is equally as competitive as WT in nasopharyngeal colonization (Naso) or lung infection (Lung).

### Genetic interaction screen of double mutant libraries identifies functionally redundant surface proteins

In order to identify functionally redundant pairs within the selected 28 non-essential surface proteins, we constructed 28 libraries, which collectively contained all 378 pairwise combinations of double mutants. Each of the 28 individual libraries contained one primary deletion in combination with 27 different secondary deletions (Fig. 1). Using this strategy, each double mutant is constructed and tested twice independently. For example, when making double mutants ΔSP_0057 ΔSP_0117 and ΔSP_0117 ΔSP_0057, in the former mutant SP_0057 was locus 1 and SP_0117 was locus 2, while in the latter mutant the reverse is true. Each double mutant library was screened for fitness in two assays; 1) colonization of the mouse nasopharynx, and 2) lung infection in mice. Genomic DNA was isolated from each inoculum and from *S. pneumoniae* that were collected from the lung and nasopharynx of challenged mice. The genomic junctions of each secondary gene deletion were amplified by PCR (Fig. 1), and each sample was individually barcoded. The frequency of each mutant in the input and output populations was subsequently determined using high throughput sequencing and the resulting read counts were used to calculate a competitive index for each mutant (see Methods).

**Figure 1.**
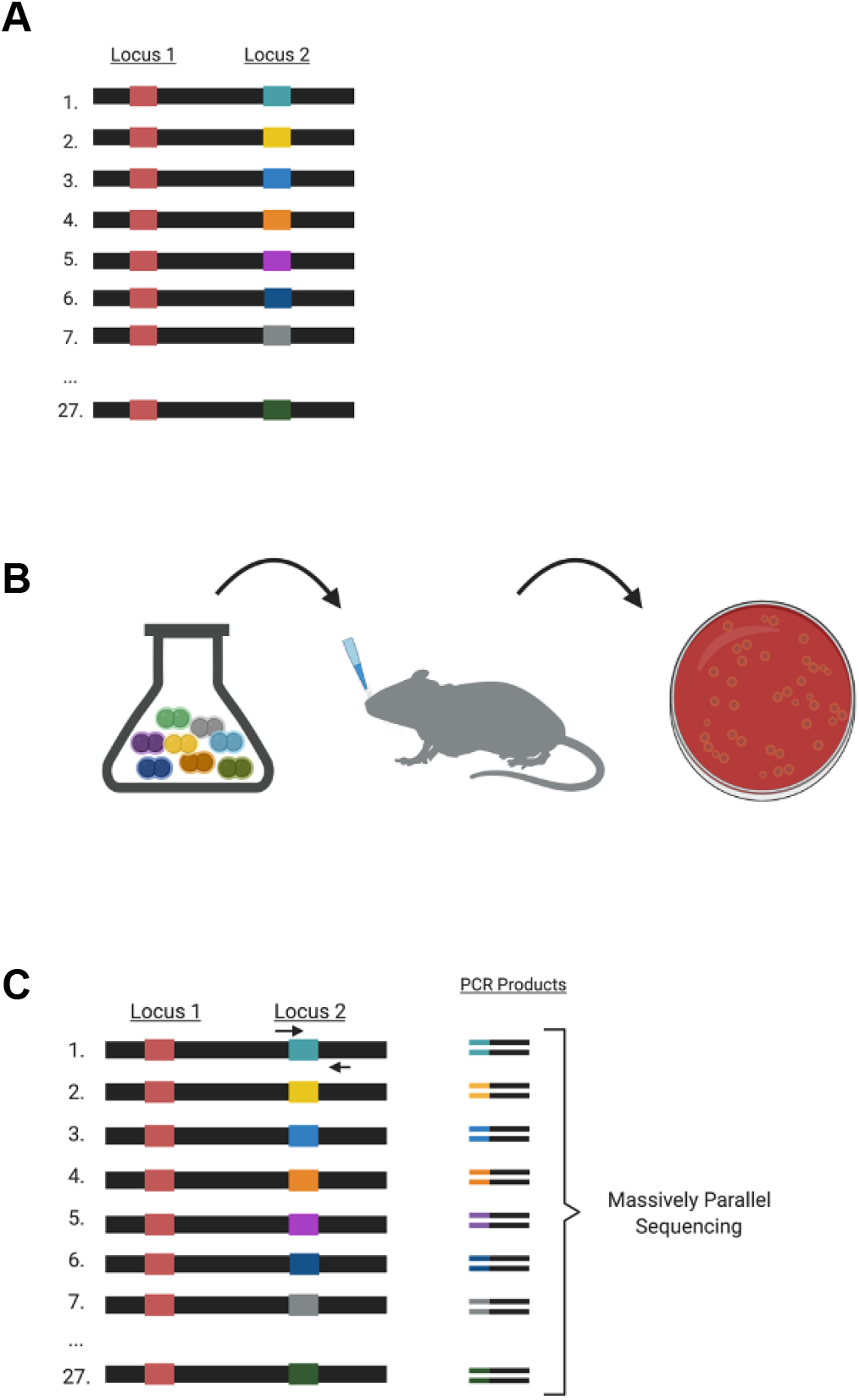
Screening double mutant libraries. (**A**) Natural transformation with pooled genomic DNA was used to create 28 double mutant libraries. One example library is shown. Within the library, every genome contains the same marked primary gene deletion (locus 1) and a varied marked secondary gene deletion (locus 2). (**B**) Each double mutant library was grown in broth culture and inoculated into groups of three mice per library, then subsequently collected from the lung and nasopharynx and outgrown on blood agar. Not all 28 mutants are depicted in the flask. (**C**) PCR strategy to determine the relative frequency of each double mutant. Locus 2 junctions were PCR amplified using a forward primer specific to the antibiotic marker in locus 2 and a tagmentation-specific reverse primer. High throughput sequencing of the PCR products was used to determine the fitness of each double mutant. (Created with Biorender.com)

All double mutant pairs were present in the input libraries, suggesting that none are synthetically lethal in vitro. From the screen, four gene pairs displayed a colonization defect in the nasopharynx and/or a defect in lung infection (Fig. 2). There was good agreement in competitive index values for most of the double mutant duplicates, with the exception of the Δ*SP_0667* Δ*ply* and Δ*ply* Δ*SP_0667* strains, which exhibited substantial variation in the nasopharyngeal colonization model, and Δ*cbpI* Δ*zmpC* and Δ*zmpC* Δ*cbpI* strains, which exhibited substantial variation in the lung infection model. For three of the four gene pairs the fitness of each single deletion was reported to be neutral in both the lung and nasopharynx. (Table 1) (22), indicating that the genes exhibit a synergistic genetic interaction. The fourth gene pair (SP_0667 SP_1923) has a synergistic genetic interaction during lung infection, since the single mutants were reported to be neutral in that infection model (Table 1) (22). However, there is no evidence of a synergistic genetic interaction in the nasopharynx colonization model, since each single mutant was reported to be attenuated in that model (Table 1) (22). We selected the most severely attenuated synergistic pair, Δ*cbpC* (choline binding protein C; SP_0377) Δ*cbpJ* (choline binding protein J; SP_0378), for further study.

**Figure 2.**
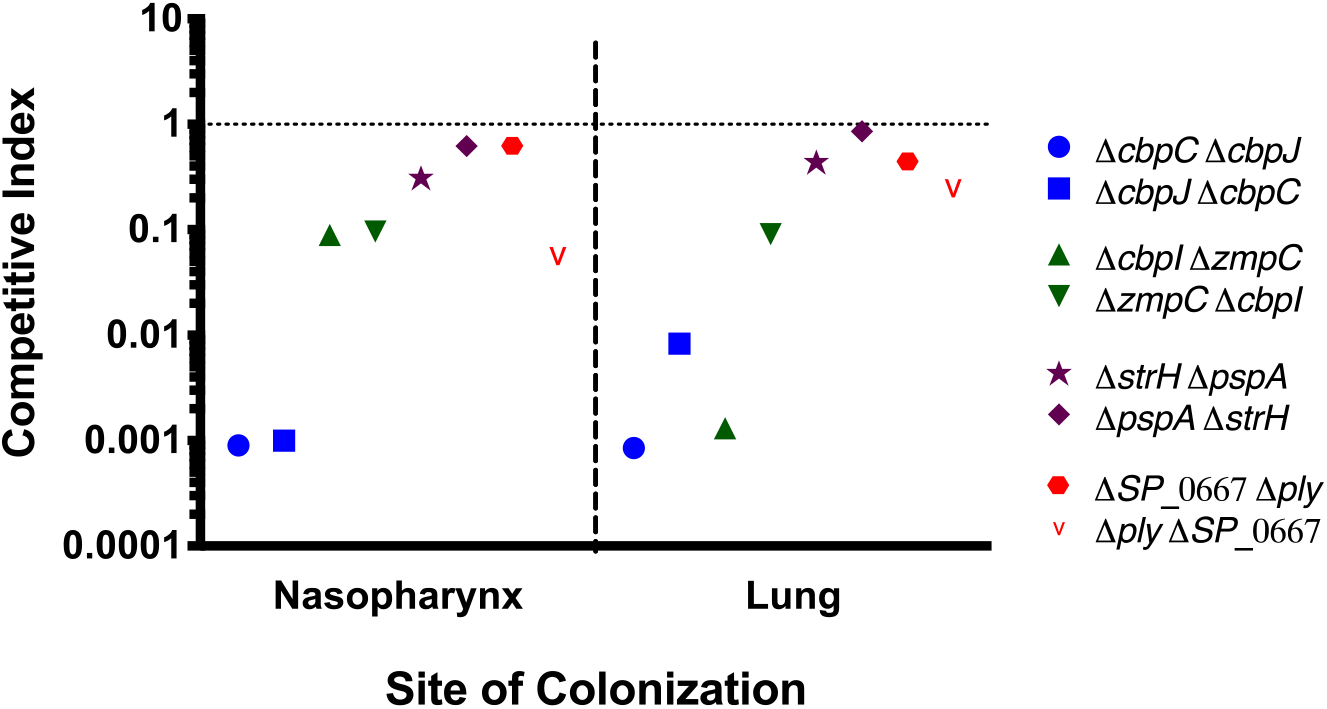
Significantly attenuated gene pairs. The competitive index (CI) was calculated for each double mutant in the library after challenge with nasopharyngeal colonization and lung infection. CI is calculated as the normalized number of reads in the output divided by those in the input. Each double mutant is represented twice throughout the double mutant libraries. The first gene listed is the primary gene deletion, followed by the secondary deletion. Significance was determined using a Student’s t-test with Bonferroni correction for multiple comparisons and all of the gene pairs displayed met a significance threshold of 0.00185.

To confirm the results of the genetic interaction screen, we reconstructed the double mutant (Δ*cbpC* Δ*cbpJ*) along with each single mutant through deletion and replacement of the coding sequence of each gene with an antibiotic-resistance cassette. Conditions of the screen were replicated by performing competitions of the mutant strains against wild-type. These experiments recapitulated the results of the screen, confirming that each single mutant has a neutral or near neutral fitness, while the double mutant is attenuated in colonization and in particular for lung infection (Fig. 3). Conversely, both single and double mutants have indistinguishable growth rates from wild-type when grown in THY broth (Fig. 3C). Taken together, these results demonstrate that *cbpC* and *cbpJ* have a synergistic genetic interaction during pneumococcal colonization and infection, but not in broth culture, suggesting that these proteins perform functionally redundant roles as virulence factors and contribute to the pathogenesis of the bacterium.

**Figure 3.**
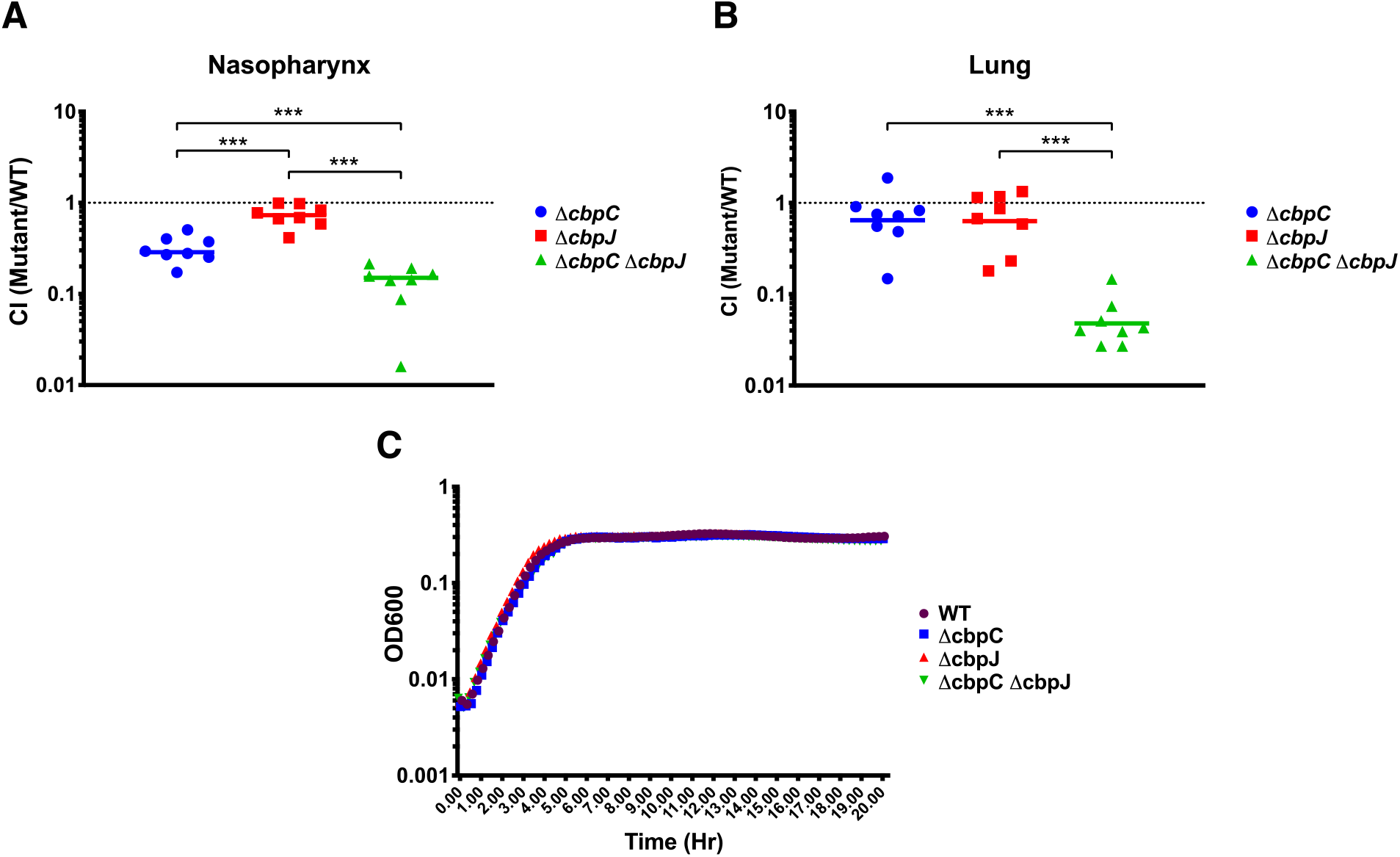
Competition and growth curve of attenuated pair Δ*cbpC* Δ*cbpJ.* *S. pneumoniae* mutant strains Δ*cbpC* (ΔSP_0377), Δ*cbpJ* (ΔSP_0378), and double mutant ΔSP_0377 ΔSP_0378 were competed against wild-type in mouse colonization (**A**) and lung infection (**B**). Competitive indices represent the output ratio (mutant to wild-type) divided by the input ratio. Each data point represents a single mouse. Statistical analyses were performed using a non-parametric Mann-Whitney U two sample rank test (***, P≥0.001). A competitive defect below that of either single mutant was observed in the double mutant during both colonization and lung infection. Bacterial growth was also analyzed by plate reader (**C**) and demonstrated similar growth kinetics for all strains when grown in THY.

### Evaluation of bivalent protein vaccine efficacy in a mouse model of pneumococcal pneumonia

In order to assess how protective these proteins are as mono- or bivalent vaccines, CbpC and CbpJ absent their signal peptides were expressed in *E. coli* and purified (Fig. S2). Mice were vaccinated with each protein alone, in combination, or mock vaccinated according to the schedule shown in Figure 4. Two weeks following the second boost, mice were challenged with *S. pneumoniae* lung infection. The median survival of the mock vaccinated mice was 1.8 days, while in the dual protein vaccinated group the median survival was extended to 2.7 days, demonstrating that vaccination with the bivalent protein vaccine resulted in a significant delay in time to death (Fig. 4B). The Prevnar vaccinated mice showed complete protection, as expected. Immunization with CbpC and CbpJ induced the production of high titers of specific antibodies against the vaccine proteins in the serum of immunized mice, as compared to mock immunized serum (Fig. 4C). Some cross-reactivity was observed between the sera of mice vaccinated against one choline binding protein in the ELISA for the other closely related choline binding protein. This is likely due to the 49% identity between CbpC and CbpJ in TIGR4. These results demonstrate that vaccination with the bivalent protein vaccine resulted in the production of specific antibodies which lead to longer survival times when challenged with pneumococcal pneumonia.

**Figure 4.**
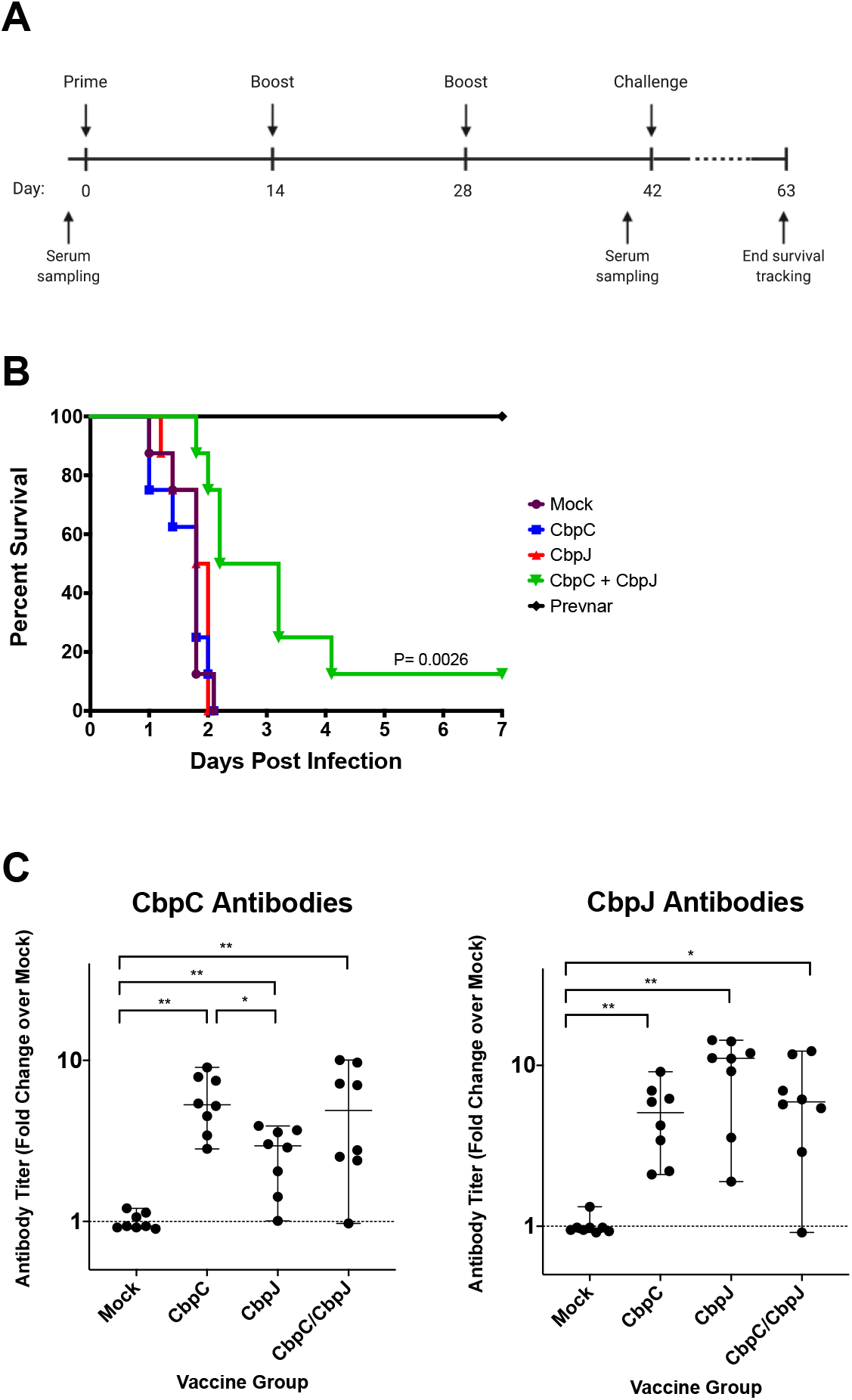
Survival of mice after challenge with Pneumococcal pneumonia. (**A**) Groups of 8 mice were immunized by intraperitoneal injection with a 1:1 mixture of protein (20 μg) to alum adjuvant. Groups were vaccinated three times at two-week intervals with CbpC, CbpJ, CbpC + CbpJ, Prevnar, or PBS (Mock). Serum was collected one day before the primary immunization and then on day 41 prior to pneumococcal challenge. (**B**) Immunized mice were challenged intranasally with 1.5×10^7 CFU TIGR4 at day 42 and then tracked for survival. Mice immunized with the divalent vaccine showed a significant delay in time to death (P=0.0026, log-rank test) and Prevnar vaccinated mice were protected as expected. (**C**) Mouse serum from day 41 (post-vaccination) was tested for specific antibodies against the protein vaccine antigens. ELISA data is displayed as the median fold change of absorbance values for vaccinated serum over mock vaccinated serum with 95% confidence intervals. Vaccination with each protein resulted in the production of specific antibodies against that protein in addition to some cross-reactivity with the other choline binding protein. ELISA data was analyzed using a Wilcoxon rank sum test (*, P≥0.033; **, P≥0.002).

## Discussion

Vaccines against *S. pneumoniae* are limited in their scope by their focus on capsular polysaccharides and as such, immunization is only able to protect against serotypes that are represented in the vaccine. The strong selective pressure of widespread vaccination with serotype-based vaccines encourages the emergence of new serotypes as disease causing strains in addition to capsular switching between strains (14–21). Therefore, there is a need for a pneumococcal vaccine that targets one or more well-conserved bacterial proteins that are essential for virulence and consequently will limit the appearance of escape mutants in response to vaccination. Because most surface proteins in *S. pneumoniae* appear to be dispensable for virulence, we sought to expand the number of potential vaccine targets. Here we present a high throughput method to identify functionally redundant gene pairs that together are essential for virulence and have the potential to limit immune escape following widespread vaccination.

To identify redundant surface proteins, we took advantage of a Tn-seq screen that was previously done in our lab (22). We selected a set of surface proteins whose disruption by transposon insertion resulted in a neutral or near neutral fitness during colonization and lung infection, which opened up the possibility that some of them may be functionally redundant. Each double mutant combination was created by making double mutant libraries, and these libraries were screened for fitness defects. This genetic interaction screen was able to identify four gene pairs that were attenuated during infection and colonization. No pairs were synthetically lethal or had a significant defect during in vitro growth, suggesting that none of the pairs perform a function that is essential for viability or aspects of basic growth and division. A fitness defect was only detected during mouse colonization and lung infection, which suggests that these proteins are dedicated virulence factors. This is a useful quality in a vaccine antigen because these proteins are expressed during infection and therefore are available to be targeted by the immune system.

The most severely attenuated pair we found was a double mutant of two choline binding protein genes, *cbpC* and *cbpJ*. Choline binding proteins are surface associated by virtue of a repeated motif that binds choline containing wall teichoic acids (35). They are important surface proteins that have roles in bacterial cell physiology and interactions with the host (36). *S. pneumoniae* strains have between 13-16 different choline binding proteins (37), which have a variety of functions including autolysis, cell division, and adherence to epithelial cells. The role of some choline binding proteins remains unknown. Due to their accessibility on the bacterial cell surface and their importance during pathogenesis, other choline binding proteins have been tested as vaccine antigens. *S. pneumoniae* PspA and PspC were both found to be protective as individual protein vaccine antigens (38–42). However, these two proteins exhibit a high degree of variability across different isolates due to their mosaic structure (42–46), making them less effective as broadly protective antigens.

Vaccination with CbpC and CbpJ elicited the production of high titers of specific antibody. Upon infection, those mice that had been vaccinated with the bivalent vaccine displayed a delayed time to death above either single protein or mock vaccinated groups, demonstrating a synergistic effect of vaccinating with multiple proteins. However, the bivalent vaccine was unable to increase overall survival. Thus, although the protection elicited is not strong enough to warrant their use as a pneumococcal vaccine, it may be effective if used in combination with polysaccharide-based vaccines to broaden protection. Additionally, use of a different immunostimulatory adjuvant or delivery site may improve effectiveness of the bivalent vaccine. Two additional limitations of our study are, first, we only interrogated 28 surface or secreted proteins, whereas many more exist, and second, we relied upon mouse models of colonization, infection and immunity, which may differ from those in the natural human host with respect to the *S. pneumoniae* proteins studied. Nevertheless, future work is warranted to investigate how protection elicited by these and other redundant protein pairs can be improved to develop a broad coverage vaccine.

Many choline binding proteins have been demonstrated to be important virulence factors (36), and have been the focus of intense study. One of our selected vaccine antigens, CbpJ, has been identified as a virulence factor with a multitude of functions: CbpJ suppresses neutrophil killing activity (47), interacts with human innate immunity protein CRP (48), and may impact adhesion to host cells (49). However, the role of CbpJ in adhesion remains unclear as some studies have found no difference in epithelial cell adhesion between Δ*cbpJ* and wild-type (47, 50). Yamaguchi et al. (47) demonstrated that a *cbpJ* deletion mutant is defective in lung infection; intranasal lung infection with Δ*cbpJ* resulted in decreased mortality in mice and lower bacterial load in the lungs. This deficiency was less pronounced during co-infection when *cbpJ* was competed against wild-type, suggesting that the wild-type may be able to complement the *cbpJ* mutant in trans (47). Because all of the lung infections in our study were performed as competitions, this can explain why we did not observe a virulence defect in the *cbpJ* deletion mutant. Although this raises the possibility that our single deletion mutants may not have a neutral fitness during single strain infection, the double mutants still display a synergistic genetic interaction. Therefore, this does not take away from our premise that vaccination with both proteins in combination will discourage the emergence of immune escape mutants.

The function of CbpC is less well-characterized, however antibodies against CbpC were identified in serum from convalescent patients, suggesting that this protein is both surface exposed and immunogenic in humans (32). CbpC was also recently shown to suppress degradation of intracellular *S. pneumoniae* by xenophagy (51), although the relevance of intracellular replication or persistence of *S. pneumoniae* is unclear given that intracellular growth is considered to be dispensable for virulence. As the CbpC-CbpJ pair was demonstrated to have functional redundancy during infection, it remains possible that CbpC performs similar functions to CbpJ, enabling it to function in place of CbpJ. Further research on these proteins, in particular CbpC, is required to better characterize their role during infection. CbpC was previously identified as a virulence factor in a transposon mutant screen for genes important during lung infection (23). However, the phenotype was not validated in that study. The discrepancy between that screen and our finding that a CbpC deletion mutant remains virulent, could be explained by a polar effect, since *cbpC* lies upstream of *cbpJ* in an operon. The transposon insertion in *cbpC* likely also disrupted expression of *cbpJ*, creating a pseudo double knockout.

In TIGR4, *cbpC* and *cbpJ* are present in an operon, however this is not the case in all sequenced strains of *S. pneumoniae*. When this project was initiated, *cbpC* and *cbpJ* were present in all publicly available *S. pneumoniae* genomes. However, when including recently sequenced strains, we found that only 39/74 genomes contain both genes. All but one of the remaining 35 strains contain one of the two, with 28 carrying *cbpC* alone and 6 with *cbpJ* alone. We propose that the vaccine may still be effective against strains that have only one of the genes in the operon because that single remaining gene may be essential for virulence without its redundant partner.

In this study, we have demonstrated an effective method to identify functionally redundant virulence factors in a bacterial pathogen, which can then be evaluated as bivalent vaccine antigen candidates. We propose that this approach could be applied to other bacterial pathogens to identify vaccine candidates that will be resistant to population level immune evasion in response to widespread vaccination. Future protein-based vaccine development pipelines should take immune evasion into account and focus on proteins that are essential or functionally redundant to prevent future escape of the pathogen from vaccination efforts.

## MATERIALS AND METHODS

### Bacterial strains and culture conditions

*Streptococcus pneumoniae* TIGR4 (serotype 4) was cultured statically in Todd Hewitt broth with 5% yeast extract (THY, BD Biosciences) supplemented with 300 U/ml catalase (Worthington Biochemicals) or on blood agar plates at 37°C with 5% CO_2_. All marked deletion mutants were made using allelic exchange where a PCR product produced with splicing by overlap extension (SOE) PCR (as previously described (52)) was transformed into *S. pneumoniae* and selected for on blood agar plates containing the relevant antibiotic. Antibiotics were used at the following concentrations: chloramphenicol (4 μg/m), spectinomycin (200 μg/ml).

### Construction of double mutant libraries

Each of the 28 genes of interest (primary genes, Table 1) were deleted and replaced with a chloramphenicol resistance (Cm^R^) cassette, then subsequently transformed with a pool of genomic DNA marked with spectinomycin resistance (Sp^R^) to replace the remaining 27 genes (secondary genes), producing a pool of double mutants. Genomic DNA concentrations were optimized to favor the production of double mutants over triple mutants. The resulting 28 double mutant pools were stored as 20% glycerol stocks at -80°C.

### Mouse infection with double mutant libraries

Inocula were prepared for infections by thawing glycerol stocks of double mutant libraries and diluting 1/20 into THY. Cultures were grown to mid-exponential growth phase, then bacteria were washed and resuspended in PBS at a concentration of 1.25×10^9^ CFU/ml. Samples of the input inoculum were frozen at -80°C for future DNA extraction. For lung infection challenge, 10-11 week old female Swiss Webster mice (Charles River) were anesthetized with 2.5% isofluorine, and then inoculated intranasally with 5×10^7^ CFU in a 40 μl volume. All mice in this study were housed in a specific-pathogen free facility at Tufts University, and all procedures were performed in accordance with Institutional Animal Care and Use Committee guidelines. Each group, which consisted of 3 mice, was challenged with a different double mutant library. After 36 h, mice were euthanized and bacteria were isolated from the lung and nasopharynx as previously described (53) and dilutions were plated on blood agar for overnight growth. The colonies were subsequently collected and frozen at -80°C for future DNA extraction.

For *in vitro* growth assessment, libraries prepared for mouse infection were also used to inoculate a culture of THY, which was grown to mid-exponential phase and subsequently frozen at -80 for future DNA extraction.

### Competition Assays

Marked deletion mutants (Cm^R^ or Sp^R^) were mixed 1:1 with the wild-type strain. The pneumococcal pneumonia mouse infection model and THY broth culture were performed as described above. Input and output samples were plated on blood agar with and without selective antibiotic to determine ratios of mutant to wild-type. Competitive index was calculated by dividing the mutant/wild type output ratio by the input ratio.

### Sequencing and analysis of double mutant libraries

Genomic DNA was isolated from input and output double mutant libraries using the DNeasy Blood and Tissue kit (Qiagen). Genomic junctions of each secondary gene replacement were amplified by first Tagmenting the DNA with TDE1 from the Nextera DNA Sample Preparation Kit (Illumina) and subsequently performing PCR with primers specific for the Nextera transposon (2A-R, Nextera) and Sp^R^ gene (5’-CTTGGAGAGAATATTGAATGGACTAATGAAAATG-3’). Each sample was uniquely barcoded using a nested PCR. Sequencing was performed as 50-bp single-end reads on an Illumina HiSeq 2500 at the Tufts University Core Facility.

The resulting sequencing data was analyzed by mapping reads to a database containing 60 bases of sequence flanking each secondary gene using CLC Genomics Workbench (Qiagen). Total read counts at each locus were normalized to the secondary gene with the greatest number of reads in each sample. Competitive index was determined by comparing the ratio of each secondary gene in the output to the ratio present in the input. Competitive indices were determined for lung infection, nasopharyngeal colonization, and in vitro growth in THY. Statistical significance was determined using a Student’s t-test with Bonferroni correction for multiple comparisons.

### Cloning, expression, and purification of recombinant proteins

Pneumococcal proteins were cloned without their signal peptide (amino acid 28 on) from strain TIGR4. These fragments were ligated into N-terminal 6xHis-tag containing vector pQE-30. Recombinant protein was expressed by IPTG induction in XL1 Blue *Escherichia coli* from the pQE-30 vector. Protein was purified using Ni-NTA agarose beads according to the manufacturer’s instructions (Qiagen). A buffer exchange was performed by centrifugation of the protein preparation through an Amicon Ultra-15 centrifugal filter unit (Millipore) with a 10 KDa molecular weight cutoff and subsequent resuspension in PBS. Proteins were purified to ≥ 90% purity as assessed by SDS-PAGE and PageBlue protein staining (Thermo Scientific).

### Mouse vaccination and challenge

Groups of 8, 10-11 week old Swiss Webster female mice (Charles River) were vaccinated intraperitoneally three times at two week intervals. Mice received 20 μg protein antigen (vaccinated) or PBS (mock-vaccinated) mixed 1:1 with Imject Alum adjuvant (Thermofisher Scientific) in a total volume of 100 μl. As a positive control, one group of mice was vaccinated intraperitoneally with 0.22 µg Prevnar per mouse, and then boosted on the same schedule as the other groups. Serum was collected via tail vein bleed prior to vaccination and one day before challenge.

At day 42, mice were challenged intranasally with 1.5×10^7^ CFU of *S. pneumoniae* TIGR4 as described above. Survival was tracked for 21 days post infection, and the survival time for each mouse was recorded.

### Endpoint ELISA

Specific antibody titers were determined by enzyme-linked immunosorbent assay (ELISA). Purified proteins were deposited into a Nunc Maxisorp 96-well microtiter plate at 5 μg/ml in 0.1 M carbonate, and incubated at 4° C overnight. Plates were washed 5 times using phosphate buffered saline with 0.05% Tween 20 (PBST), and then blocked in PBST with 3% BSA for 1 hour at room temp. All subsequent washes were performed using PBST with 0.2% BSA. Plates were then washed 5 times and serum samples were diluted 100-fold into the first well and then serially diluted 20-fold for all subsequent wells and incubated at 4° C overnight. Plates were then washed 5 times and incubated with goat anti-mouse Ig conjugated to HRP (SouthernBiotech) for 1 hour at room temp. Plates were washed again 5 times and developed with KPL SureBlue TMB 1-component peroxidase substrate (SeraCare). Following 30 minutes of incubation, color development was stopped with KPL TMB BlueStop solution (SeraCare) and absorbance of plates was read on a plate reader at 650 nm. The endpoint antibody titer is defined as the fold change of absorbance of each vaccinated serum sample over mock vaccinated serum.

## ACKNOWLEDGEMENTS

The authors would like to thank Ognjen Sekulovic for insightful scientific discussion, as well as Tufts University Core Facility, Albert Tai, and Tufts Comparative Medicine Services for experimental assistance. This research was supported by National Institutes of Health grants AI110826 (A.C.) and training grant GM139772 (A.J.M.).

## AUTHOR CONTRIBUTIONS

**LMR:** Investigation, Data Curation, Methodology, Writing - Review & Editing **AJM:** Writing - Original Draft Preparation, Investigation, Data curation, Methodology **RFM:** Investigation, Methodology, Writing - Review & Editing **DL:** Methodology **AC:** Conceptualization, Funding acquisition, Writing - Reviewing & Editing

